# A Versatile AAV-TH-SNCA Model to Study Early α-Synuclein Pathology and Intervention

**DOI:** 10.64898/2026.03.20.712963

**Authors:** Sabina Marciano, Leandra G. Velazquez, Claudia Rodríguez López, Neranjan de Silva, Garrett Sommer, William Tower, Eileen Ruth S. Torres, Michael G. Kaplitt, Teresa A. Milner, Roberta Marongiu

## Abstract

Adeno-associated viral (AAV)-mediated overexpression of human wildtype α-synuclein (α-syn) in the substantia nigra (SN) is a widely used approach to model Parkinson’s disease (PD) in rodents. However, variability in the ability of AAV-based systems to induce nigrostriatal pathology and motor deficits has limited reproducibility across studies, especially in mice. Here, we systematically optimized key vector features - AAV serotype, promoter, viral titer - to establish a highly efficient and reliable mouse model of PD. We compared the tropism and expression efficiency of mixed AAV2/1 and AAV2/rh10 serotypes combined with three promoters - CMV enhancer/chicken β-actin (CBA), human Synapsin (hSYN), and rat Tyrosine Hydroxylase (TH) - to drive human α-syn gene (SNCA) expression in nigral dopaminergic neurons. The AAV.TH.SNCA vector, delivered at an optimized titer, achieved selective and sustained α-syn overexpression in nigral neurons, resulting in nigro-striatal neurochemical changes and progressive motor deficits preceding overt neuronal loss. Fine tuning α-syn expression proved critical for detecting early disease processes: lower AAV.TH.SNCA titer induced early pathological signatures, including α-syn hyperphosphorylation and neuroinflammation, whereas higher titers produced robust nigrostriatal degeneration not achieved with other promoter constructs. Notably, we demonstrate that motor and neurochemical impairments can occur prior to dopaminergic cell death, implicating microglial activation and α-syn pathology as primary drivers of dysfunction. This observation is consistent with human genetic evidence showing that triplication of the wild-type SNCA gene alone can cause Parkinsonian pathology, highlighting that our model enables the use of a single experimental reagent to investigate the molecular, cellular, and behavioral consequences of controlled increases in α-syn expression. This novel AAV.TH.SNCA model provides a powerful and versatile platform for investigating mechanisms of a α-syn-mediated neurotoxicity and for evaluating disease modifying interventions targeting early, pre-degenerative stages of PD.

**Highlights:**

- Titrated α-syn expression uncouples early dysfunction from dopaminergic neuron loss
- AAV2/rh10-TH-SNCA model captures prodromal and degenerative PD stages
- Motor deficits arise from α-syn pathology and nigral molecular changes before neurodegeneration

## INTRODUCTION

Parkinson’s Disease (PD) is characterized mainly by the progressive degeneration of nigrostriatal dopaminergic (DA) neurons projecting from the substantia nigra pars compacta (SN) to the caudate and putamen, also named dorsal striatum in rodents^1^. SN neurons are particularly vulnerable to oxidative stress, mitochondrial dysfunction, neuroinflammation, and accumulation of misfolded/aggregated proteins ^2^. Although the underlying causes of their selective vulnerability are not completely understood, accumulating evidence suggests that early molecular changes in the SN play a critical role during the prodromal phase of PD. Moreover, preclinical and clinical studies suggest that compensatory neurochemical changes occur prior to significant neuronal loss (>50% total SN neurons) typical of more advanced disease stages ^3–6^. Among the earliest pathological features are α-synuclein (α-syn) protein accumulation, α-syn phosphorylation at Ser129 (pSer129), and microglial activation, which precede overt neurodegeneration and motor symptoms ^7–20^.

Adeno-associated viral (AAV) vectors driving expression of human wild-type α-syn in the SN are a widely used tool to model PD dysfunction in the basal ganglia in rodents. AAV features such as the serotype, promoter, and viral titer offer cell specificity and versatility, and they could be used to mimic PD stages by tuning the α-syn expression levels. Fine-tuning α-syn expression is crucial to accurately model progression of disease pathology and phenotype and elucidate mechanisms underlying regional vulnerability of DA neurons. Previous studies have employed different AAV serotypes - including AAV2, AAV2/1, AAV2/7, AAV2/9 - to induce α-syn overexpression within the rodent nigrostriatal circuit generating variable results across studies, mostly in black C57BL6 wild-type mice and Sprague Dawley rats (**Table 1**) ^9,13,15,16,18,21–30^; for more ample review ^31,32^. Most studies used high AAV titers to achieve nigral degeneration and striatal dopaminergic fiber loss, although these may result in RNA and protein expression at levels that can cause toxicity unrelated to α-syn expression. Injecting AAV.hSNCA at high titer also poses the difficulty of distinguishing early vs late PD-related phenotypes, as α-syn expression levels were not modulated across a defined range. Interestingly, a study by Oliveras-Salva et al., using AAV2/7.hSYN.SNCA, reported titer-dependent TH+ nigral cell loss at 4- and 8-weeks post-injection, followed only later by motor deficits at 12-15 weeks post-AAV injection ^18^, and work by Dr. Björklund and collaborators shows progressive nigral neurodegeneration when AAV vectors are used at high titer ^31^.

**Table 1.**
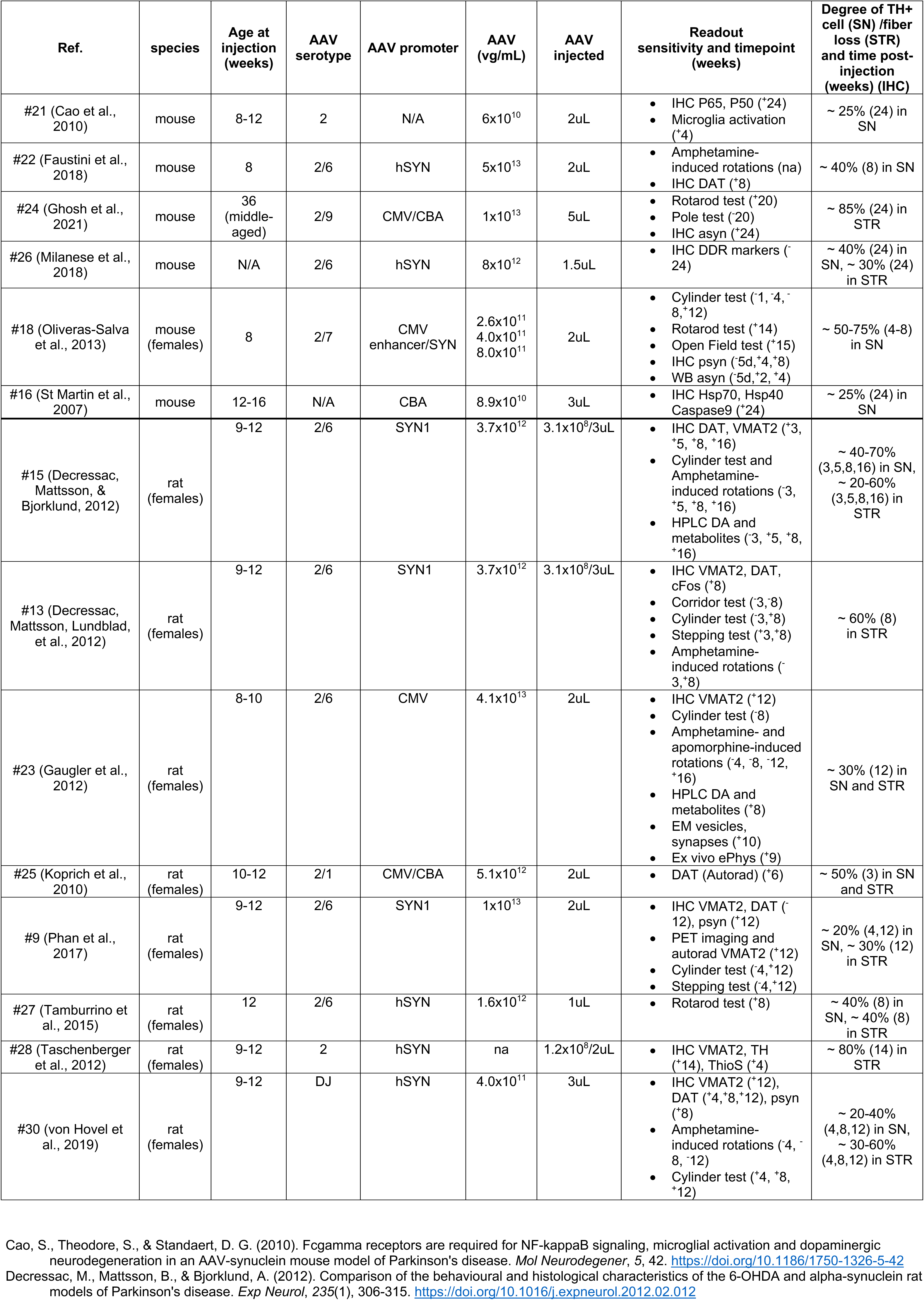
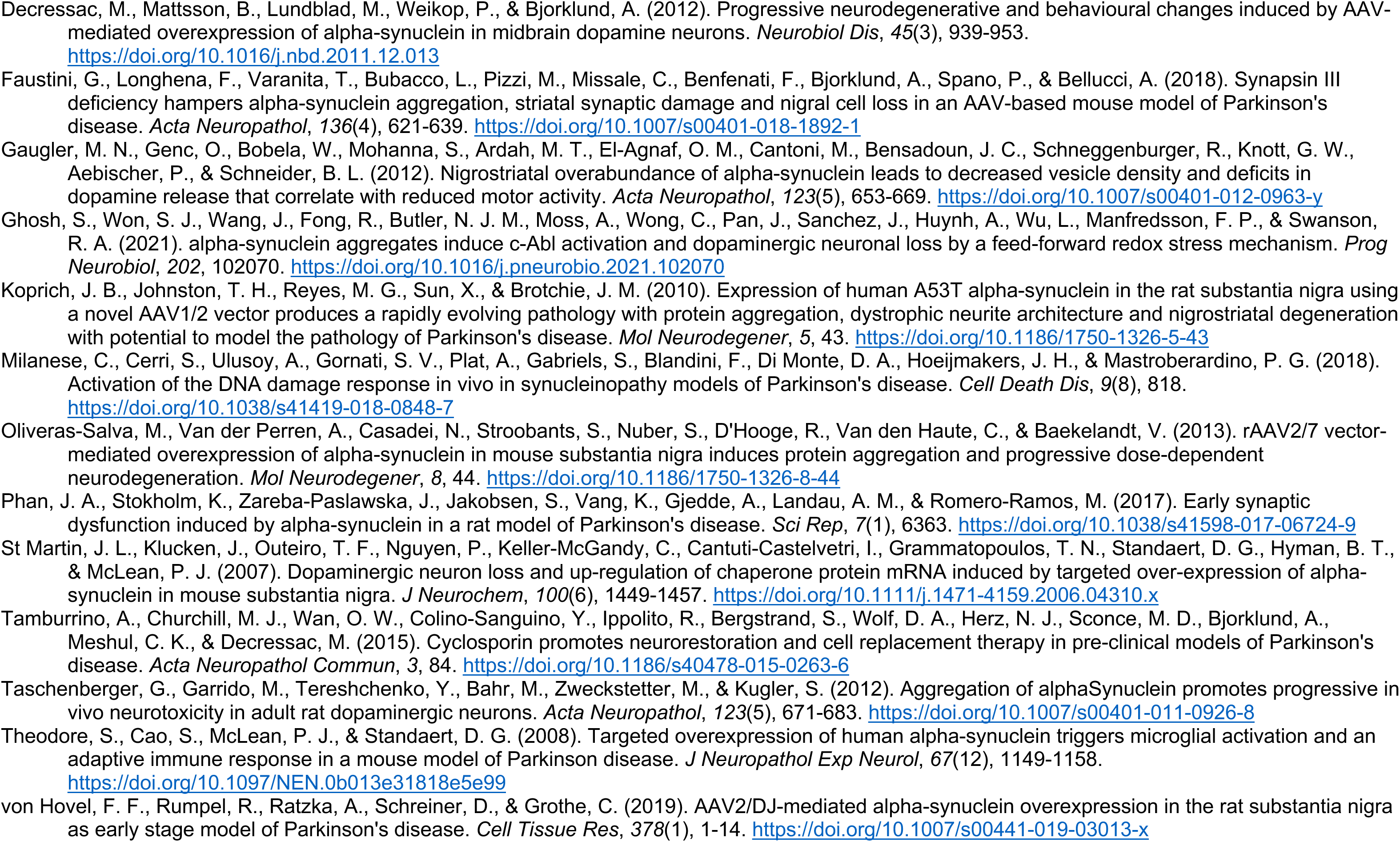
AAV-hSNCA overexpression in the substantia nigra of wild type mice or rats, and timeline of molecular changes and behavioral phenotypes. Only publications that reported information on TH loss were included in the table. Please note that information relative to AAV titers is reported as found in the methods section of these publications, which might not have used consistent descriptive metrics. CMV, cytomegalovirus; CBA, hybrid CMV enhancer/chicken β-actin promoter; hSYN, human Synapsin promoter; IHC, immunohistochemistry; WB, Western Blot; ICAM, Intercellular adhesion molecule; IL, Interleukin; DDR, DNA damage response; DAT, Dopamine transporter; VMAT2, Vesicular Monoamine Transporter 2; TH, Tyrosine Hydroxylase, SN, Substantia Nigra: STR, striatum.

These findings suggest that features such as AAV serotype, titer, and promoter may be leveraged to generate optimized AAV.SNCA vectors, enabling control of hSNCA expression to modulate key features of PD progression in animal models. However, the field has lacked a rigorously validated, titratable viral-based platform - and a systematic understanding of how promoter choice, viral serotype, and vector titer interact to regulate α-synuclein expression levels and pathology - thereby limiting investigation of early, pre-degenerative disease mechanisms and therapeutic windows.

In this study, we used different AAV vector combinations carrying the hSNCA transgene to titer the overexpression of human wild-type α-syn in the mouse SN, and longitudinally monitored neurochemical and phenotypical changes for up to eight months. We compared AAV vectors based on two serotypes - AAV2/1 and AAV2/rh10 - and three promoters - hybrid CMV enhancer/chicken β-actin (CBA), human Synapsin (hSYN), and rat Tyrosine Hydroxylase (TH) - to determine their tropism, transduction efficiency, and ability to generate a PD model with motor phenotype and nigral neurodegeneration. By systematically manipulating AAV serotype, promoter, and titer, this study provides a rigorous and quantitative framework to optimize AAV-hSNCA models, enhancing their reproducibility and translational relevance for investigating PD mechanisms.

## RESULTS

### Evaluation of AAV2/1 and AAV2/rh10 serotypes and CBA, hSYN, and TH promoters for efficient mCherry expression in nigral neurons

To optimize transgene delivery to nigral dopaminergic neurons, we first compared the efficiency of two AAV serotypes (AAV2/1 and AAV2/rh10) combined with three promoters (CBA, hSYN, and TH) using the fluorescent reporter mCherry (**Figure 1a-c**). This experiment was designed to identify potential differences in serotype-promoter combination that could influence transgene expression levels and to identify the optimal serotype-promoter combination for robust and selective transduction of SN neurons prior to introducing the human SNCA transgene.

**Figure 1.**
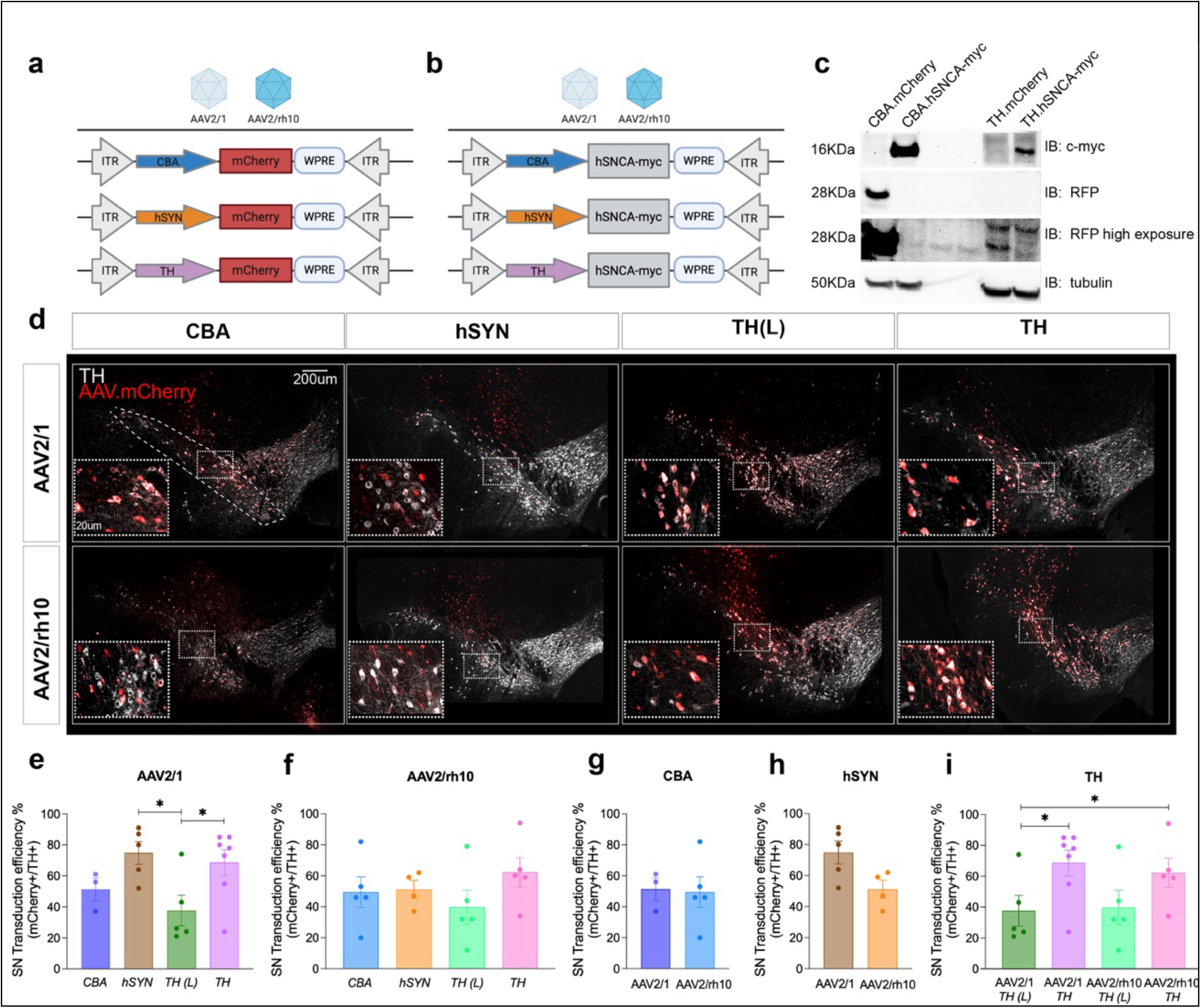
AAVs constructs, *in vitro* validation and transduction efficiency. Vectors with AAV2/1 and AAV2/rh10 serotype express mCherry (a) or human wild-type α-syn transgene (SNCA) (b) under the CBA, hSYN, and TH promoters. c. SNCA expression from pAAV plasmids in HEK-293T cells. d. Representative images of TH and mCherry staining in the SN. A layout of the region drawn in the SN for the quantification is reported in the first panel. e-i. Quantification of the mCherry+ cells normalized by total TH+ cells per area (mm^2^) shows that the transduction efficiency is higher for AAV2/1.hSYN.mCherry and AAV2/1.TH.mCherry (i). Scale bar, 200 mm. *p < 0.05, One-way ANOVA, Tukey’s multiple comparison test (F(3, 16)= 3.68, *p= 0.03). CBA, hybrid CMV enhancer/chicken β-actin promoter; hSYN, human Synapsin promoter; TH, Tyrosine Hydroxylase promoter; SN, substantia nigra; mpi, months post-injection.

The hybrid CMV enhancer/chicken β-actin promoter is a widely used strong ubiquitous promoter that drives high, but non-specific expression across multiple cells types ^33^. In contrast, the hSYN promoter restricts the expression primarily to neurons ^34^, although it typically produces more moderate expression levels. Of the three selected promoters, the rat TH promoter provides the highest degree of specificity for dopaminergic neurons, enabling targeted expression within the nigrostriatal system ^35^. The hybrid serotypes AAV2/1 and AAV2/rh10 were selected based on their known neuronal tropism and strong transgene expression in the central nervous system (CNS) ^33,36,37^. AAV2/1, although it possesses a slight lower transduction efficiency compared to other serotypes, combines the high expression efficiency from of AAV1 with the high neuronal specificity and localized spread of AAV2, which is advantageous when targeting small brain structures such as the substantia nigra. The AAV2/rh10 was selected for its high neuronal specificity (of AAV2 and AAVrh10), and high transduction efficiency per cell and relatively low immunogenicity of AAVrh10 serotype, properties that have contributed to its use in CNS-directed AAV gene therapy human studies ^38,39^.

Because previous studies have reported that AAV-mediated expression of green fluorescent protein (GFP) in the brain parenchyma can elicit neuroinflammation and DA neuron loss ^40–43^, we used AAV-mCherry vectors as matched controls for each construct to isolate the specific contribution of α-syn expression to neurodegeneration.

All AAV vectors were injected at a titer of 1×10^12^ vg/ml. Vectors containing the TH promoter were also injected at lower titer of 5×10^11^ vg/ml (designated TH(L)).

Fluorescent co-immunolabelling for mCherry and TH (to label DA neurons) revealed that both AAV serotypes produced strong mCherry expression in approximately 40–80% of TH+ neurons in the SN (**Figure 1e–i**). The highest number of TH+ neurons was transduced by the AAV2/1.hSYN.mCherry vector (74.8 ± 0.81%) and AAV2/1.TH.mCherry (68.6 ± 0.67%). Other serotype-promoter combinations achieved transduction rates of approximately 40%. These results demonstrate that both AAV2/1 and AAV2/rh10 effectively target mouse nigral neurons, with the TH and hSYN promoters providing the most robust and neuron-specific expression (**Table 2**).

**Table 2.** Summary of results.

| AAV Serotype | AAV Promoter | Transduction Efficiency | TH Loss (SN) | TH Loss (STR) | Iba1/pSyn (SN) | Motor Deficits | hSNCA mRNA Expression |
| --- | --- | --- | --- | --- | --- | --- | --- |
| 1/2 | *TH(L)* | ~40% | – | – | – | – | NA |
| 1/2 | *TH* | ~70% | Yes | Trend | – | – | NA |
| 2/rh10 | *TH(L)* | ~40% | – | Trend | ↑<br>(significant correlation) | Yes (OF, Cylinder) | ~200% increase to mCherry control |
| 2/rh10 | *TH* | ~60% | Yes | Yes | ↑<br>(significant correlation) | Yes(OF) | ~500% increase to mCherry control |
| 1/2 | *CBA* | ~50% | – | – | – | – | NA |
| 1/2 | *hSYN* | ~80% | – | – | – | – | NA |
| 2/rh10 | *CBA* | ~50% | – | – | – | – | NA |
| 2/rh10 | *hSYN* | ~50% | – | – | – | – | NA |

### Low-titer AAV2/rh10.TH-driven α-syn overexpression produces early and sustained motor deficits

To assess the functional consequences of α-syn overexpression mediated by different AAV vectors, we performed longitudinal behavioral testing (**Figure 2**). The transduction of nigral dopaminergic neurons with AAV2/rh10.TH.SNCA(L) resulted in a 45.3 ± 6.33% decrease of mouse locomotor activity in the open field test as early as 1-month post-injection, and consistently for up to 7 1/2 -months (p<0.001) (**Figure 2a, Suppl. Fig. 1c**). Compared to the corresponding AAV-mCherry controls, these mice also showed a significant 45.7 ± 11.1% reduction of wall contacts during rearing in the cylinder test starting at two months post-injection, which progressed to a 67.7 ± 10.4% decline by six months (p<0.001; **Figure 2b**, **Suppl. Figure 1c**). In the rotarod test, AAV.SNCA-injected mice showed no significant difference in latency to fall compared to corresponding mCherry controls.

**Figure 2.**
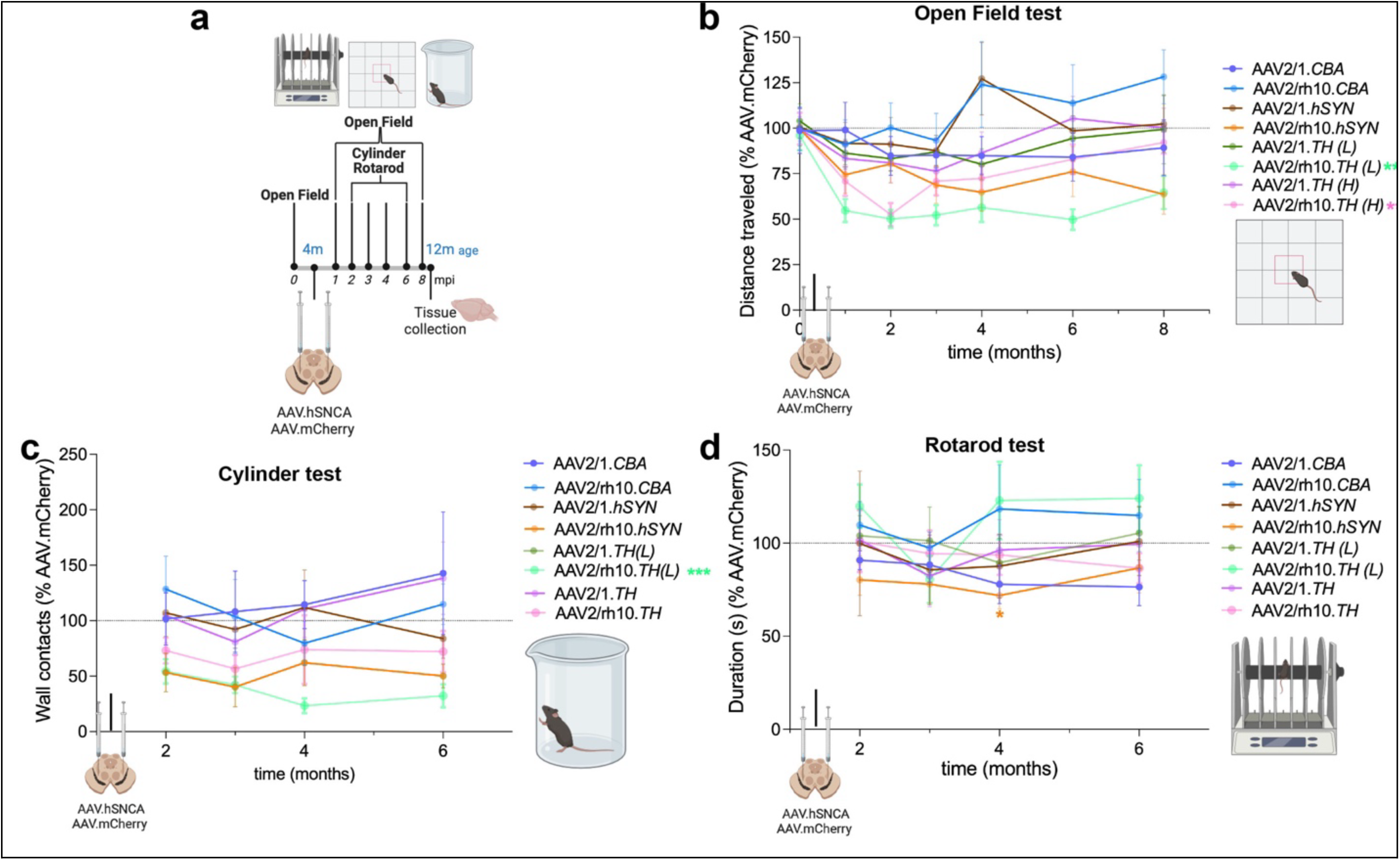
AAV2/rh10.TH.SNCA(L) induces motor deficits most efficiently than the other AAV serotype-promoter vector combinations. a. Experimental timeline for this study. b-c. Mice injected with AAV2/rh10.TH.SNCA(L) showed significant motor deficits in the open field (b) and cylinder test (c). In contrast, animals injected with other serotype/promoter AAV combinations did not display motor deficits across the behavioral tests. d. No major effects in latency to fall in the rotarod test (4-40 rpm) were observed. *p < 0.05, ***p < 0.001. Open Field, AAV2/rh10.TH.SNCA (L): Two-way ANOVA, with Sidak’s multiple comparison test, effect of time F(3.47, 41.72)= 18.42, ****p < 0.0001, effect of treatment F(1, 13)= 34.96, ****p < 0.0001, and effect of time*treatment F(3.47, 41.72)= 6.694, ***p = 0.0005. Open Field AAV2/rh10.TH.SNCA: Two-way ANOVA, with Sidak’s multiple comparison test effect of time F(3.89, 54.53)= 7.18, ****p = 0.0001, effect of treatment F(1, 14)= 18.89, ***p = 0.0007, and effect of time*treatment F(3.89, 54.53)= 7.18, ***p = 0.0001). Cylinder test, AAV2/rh10.TH.SNCA (L): Two-way ANOVA, with Sidak’s multiple comparison test effect of time F(1.58, 20.05)= 12.33, ***p = 0.0007, effect of treatment F(1, 13)= 37.38, ****p < 0.0001, and effect of time*treatment F(1.58, 20.05)= 0.104, ns p = 0.85. TH, Tyrosine Hydroxylase; CBA, hybrid CMV enhancer/chicken β-actin promoter; hSYN, human Synapsin promoter; mpi, months post-injection.

Mice injected with the higher titer AAV2/rh10.TH.SNCA vector showed significant motor deficits in the open field test (p< 0.0001) and a trend toward decrease function in the cylinder test compared to the mCherry controls. However, their motor function deficit quantified as percentage of control group was less than what observed for the low titer AAV2/rh10.TH.SNCA (L), as a small decrease in motor activity was observed over time also for the AAV.TH.mCherry control group, suggesting a slight adverse influence of the higher titer control vector on nigral neuronal function even in the absence of demonstrable cell loss (**Figure 3**).

**Figure 3.**
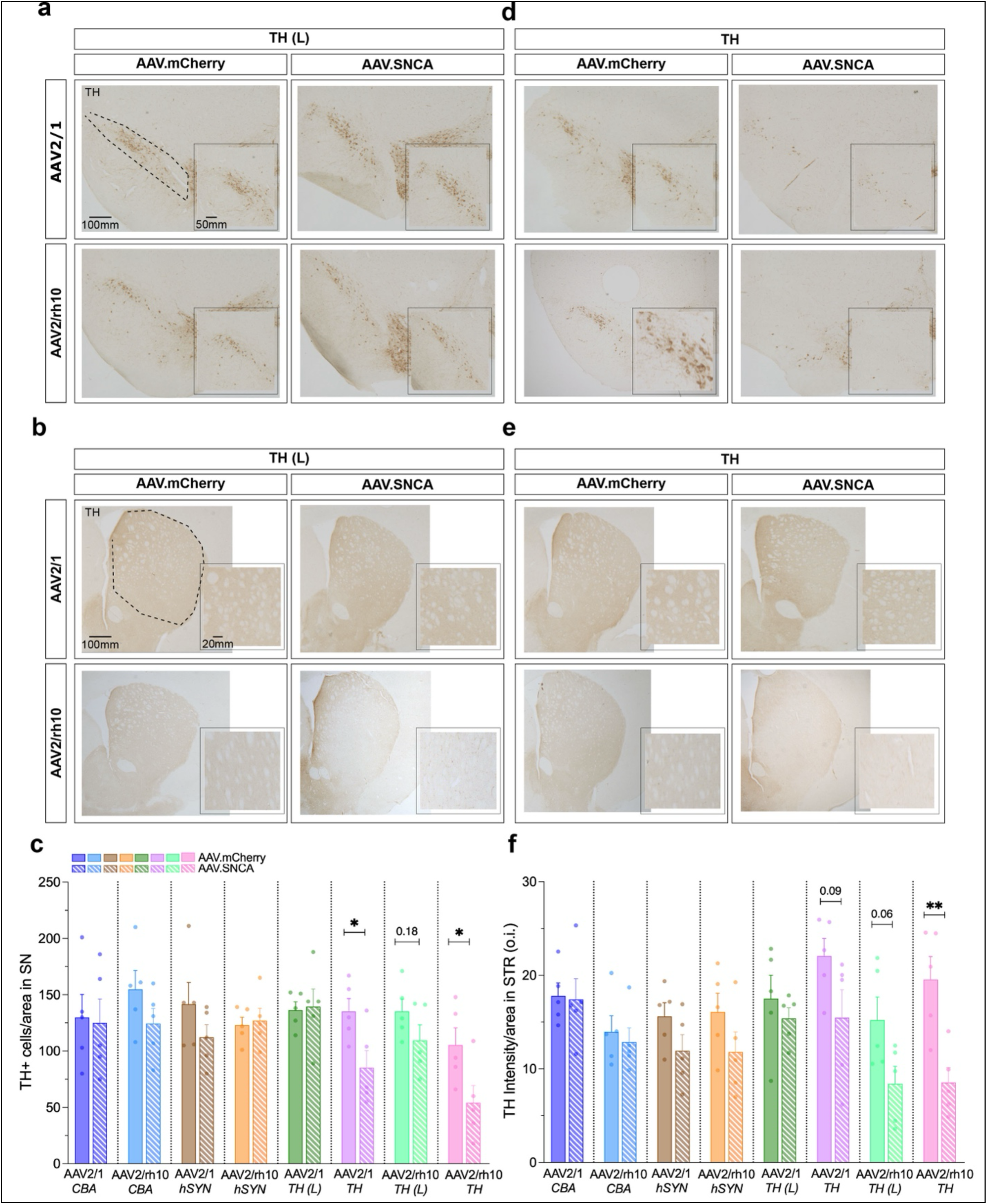
Selective neurodegeneration induced by AAV2/rh10.TH.SNCA and AAV1/2.TH.SNCA vectors following hSNCA overexpression. Representative images of TH DAB-immunolabelling in the SN (a) and dorsal striatum (b) for the AAVs.TH injected at the low (L) titer. c. Quantification of the TH+ cells per area (mm^2^) in the SN indicated no TH loss across serotype, promoter, and titers of choice in AAV.mCherry injected mice. Yet, a significant reduction was observed in the SN of mice injected only with the vectors expressing hSNCA under the rat TH promoter. d-e. Representative images of TH DAB-immunolabelling in the SN (d) and dorsal striatum (e) for the AAVs.TH injected at the high titer. f. Quantification of the TH signal intensity per selected region of interest in the striatum demonstrated no TH loss across serotype, promoter, and titers of choice in AAV.mCherry-injected mice. Yet, TH intensity was significantly reduced in the striatum of mice injected with AAV2/rh10.TH.SNCA. *p < 0.05, **p < 0.01. Unpaired t-test. TH, Tyrosine Hydroxylase; SN, substantia nigra; CBA, hybrid CMV enhancer/chicken β-actin promoter; hSYN, human Synapsin promoter.

### AAV2/rh10.TH and AAV2/1.TH vectors induce TH loss following α-syn overexpression

To determine if α-syn overexpression by our AAV vectors could induce dopaminergic neurodegeneration, following completion of behavioral testing, mice were euthanized at ∼8 months post-injection. We then assessed nigral TH expression following delivery of human wild-type α-syn or mCherry control vectors.

AAV-mCherry did not result in TH loss in either the SN and striatum, regardless of the serotype, promoter, or viral titer used (**Figure 3**). In contrast, overexpression of α-syn using the AAV2/rh10.TH vector at high titer led to a 46.9 ± 0.96% reduction in TH+ neurons compared to the corresponding AAV.mCherry control, and injection of high-titer AAV2/1.TH.SNCA caused a 37.0% ± 13.8% decrease in TH immunoreactivity in SN compared with controls (**Figure 3a-c**). Other serotype-promoter combinations did not induce significant TH level reduction at 8 months post-injection. AAV2/rh10.TH.SNCA also caused a significant decrease in TH fiber density in the striatum (AAV.SNCA 8.5 ± 3.59 vs. AAV.mCherry 19.5 ± 5.55, p=0.006); whereas a trend toward decrease was observed with injection of AAV2/1.TH.SNCA (AAV.SNCA 15.5 ± 6.73 vs. AAV.mCherry 22.06 ± 4.18, p=0.09), and no differences with the other AAV vectors expressing α-syn compared to their respective AAV-mCherry control group (**Figure 3d-f**).

These findings indicate that promoter-specific overexpression of α-syn selectively impacts dopaminergic neurons and that the TH promoter in combination with specific serotypes (particularly AAV2/rh10) produced the strongest TH neuronal loss (**Table 2**).

### Low-titer AAV2/rh10.TH.SNCA triggers α-syn hyperphosphorylation and neuroinflammation without neurodegeneration

To investigate early, pre-degenerative mechanisms of α-syn nigral accumulation, we examined the effects of low-titer AAV2/rh10.TH.SNCA (L) and AAV2/1.TH.SNCA (L) injection into the SN, which did not induce significant nigral TH loss, in comparison to the same vectors injected at higher titer and to their corresponding mCherry controls (**Figure 4**). Α-syn overexpression by AAV2/rh10.TH.SNCA (L) markedly increased the number (230.5 ± 2.74%) of Iba1+ microglial cells in the SN compared to the corresponding control group (**Figure 4a–c**). The same AAV2/rh10.TH.SNCA (L) vector induced a significant threefold increase in α-syn phosphorylation at Ser129 (333.3 ± 4.94%), a hallmark of pathological α-syn accumulation (**Figure 4a,b,d**), compared to the AAV2/rh10.TH.mCherry (L) and the higher-titer AAV2/rh10.TH.SNCA group. No TH loss was observed with AAV2/rh10.TH.SNCA (L) viral vector (**Figure 3**), suggesting that the neuroinflammatory response may be driven by pathological α-syn accumulation rather than cell death. This is supported by a significant correlation between nigral Iba+ cell number and α-syn pSer129+ cells compared to mCherry control (**Figure 4e-h**). Together, these results identify AAV2/rh10.TH.SNCA as a suitable vector for studying both early-stage and advanced, nondegenerative mechanisms of PD depending upon titer.

**Figure 4.**
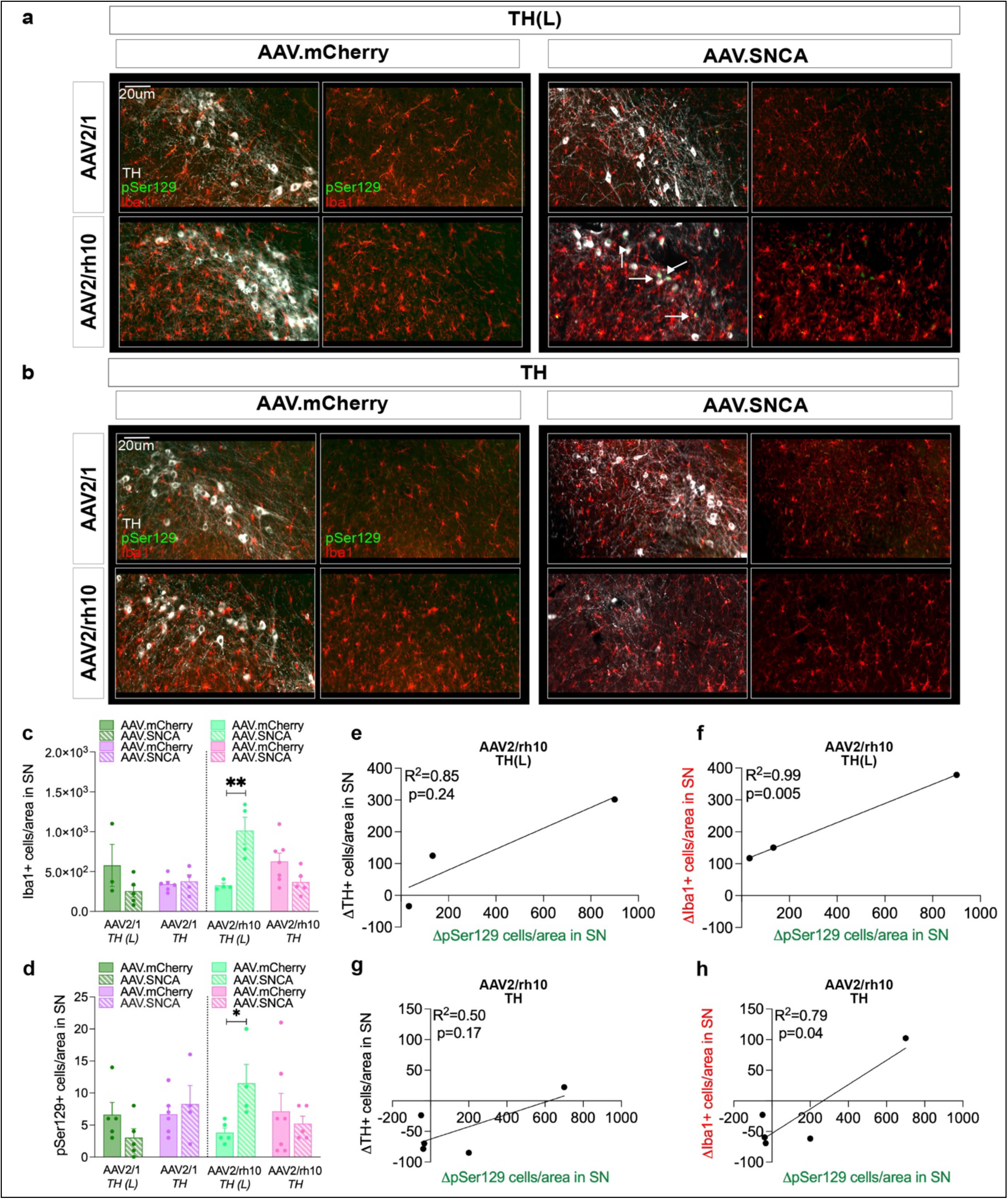
AAV2/rh10.TH.SNCA(L) triggers microglia activation and a-syn hyperphosphorylation in the SN. Representative images of Iba1 and pSer129 fluorescent immunolabelling in the SN of mice injected with AAVs.TH at low (L) (a) and high titer (b). Mice injected with the AAV2/rh10.TH.SNCA(L) showed an increase in the number of Iba1+ cells (c) and number of pSer129+ cells (d) per selected region of interest in the SN (mm^2^). (e-h) A significant correlation between the number of Iba1+ (ΔIba1+), but not TH+ neurons, and pSer129+ cells (ΔpSer129+) in the SN was found for the AAV2/rh10.TH.SNCA, quantified as percentage of corresponding mCherry controls. Simple linear regression analysis. *p < 0.05, **p < 0.01, Unpaired t-test. TH, Tyrosine Hydroxylase; SN, substantia nigra. Scale bar, 20 μm.

These results indicate that modest, TH-driven α-syn overexpression in the SN is sufficient to elicit molecular and inflammatory alterations that translate into early and persistent motor impairments, prior to observable neuronal loss, thereby providing a functional correlate of prodromal PD pathology (results summarized in **Table 2**).

### AAV titer-dependent increase in hSNCA mRNA expression in transduced nigral neurons

The differences in neurochemical and behavioral phenotype observed in mice injected with low and high titer AAV.TH.SNCA suggest that these may be mediated by differences in nigral transduction efficiency (**Figure 1j**). Graded levels of pathological α-syn accumulation in the SN due to the differential hSNCA transgene expression may also be involved. To test this, we quantified hSNCA mRNA levels using RNAscope *ISH* in nigral tissue transduced with either low or high titer AAV2/rh10.TH.SNCA vectors (n=4/group). In AAV2/rh10.TH.mCherry samples from both titer groups (n=4/group), and hSNCA transcript levels in AAV.SNCA-injected samples were normalized to the respective AAV.mCherry controls. Quantification at the single-cell level revealed a ∼two-fold increase (TH(L) 254.6 +/- 82.5 vs. TH(high) 517.6 +/- 63.8; p=0.045) in hSNCA mRNA levels in neurons transduced with the high-titer AAV.SNCA compared to the low-titer vector (**Figure 5c**). These data demonstrate that AAV2/rh10.TH.SNCA vector viral titer regulated intracellular hSNCA transcript burden per neuron at these two titers, rather than simply the number of transduced cells.

**Figure 5.**
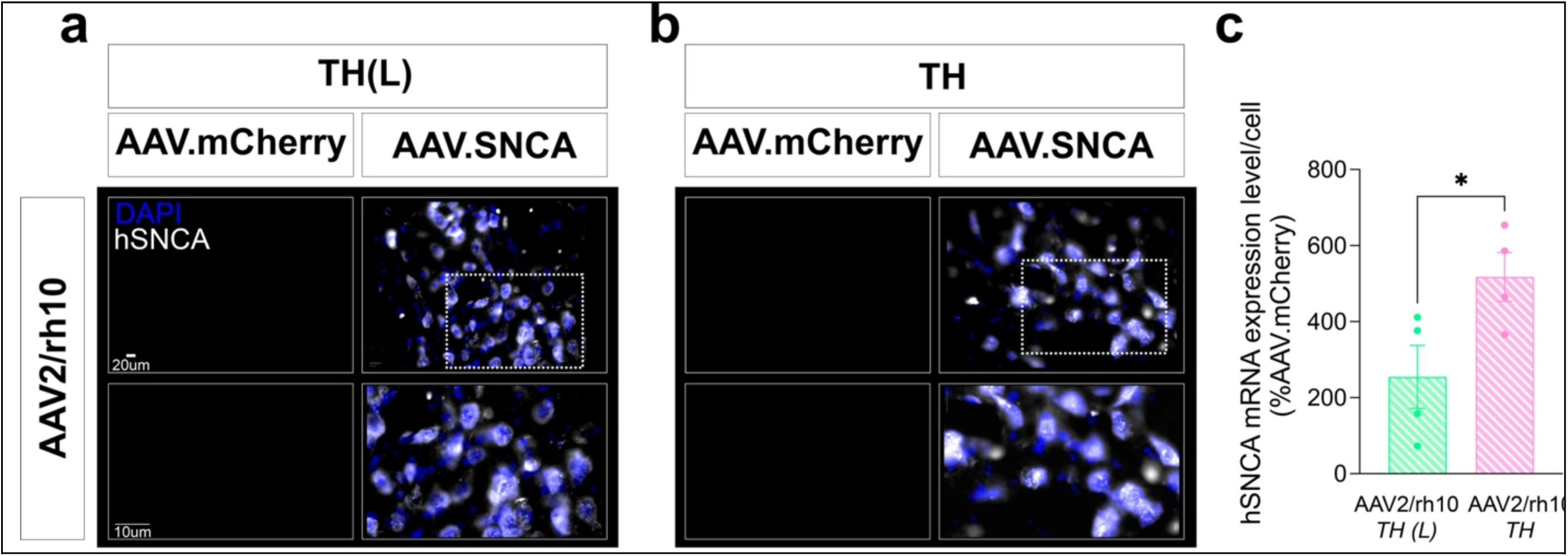
AAV titer-dependent levels of hSNCA mRNA in transduced nigral neurons. a,b. Representative images of hSNCA mRNA using RNAscope *ISH* in the SN of mice injected with either low (L) (a) or high (b) titer AAV.TH.SNCA or AAV.mCherry control. c. Quantification of hSNCA mRNA levels at single-cell resolution (percent change to respective AAV.mCherry controls) revealed a ∼two-fold increase in neurons transduced with the high titer AAV.TH.SNCA compared to the AAV.TH.SNCA (L). n=4. *p < 0.05, Unpaired t-test. Scale bars, 10 and 20 μm.

## DISCUSSION

In this study, we developed and systematically validated an optimized AAV vector mouse model of PD based on overexpression of human SNCA gene in nigral dopaminergic neurons. By directly comparing AAV2/1 and AAV2/rh10 serotype tropism for dopaminergic neurons, hSNCA transgene expression under CBA, hSYN, or TH promoter, and two viral titers, we optimized an AAV vector system that achieved selective and sustained transduction of nigral dopaminergic neurons and distinct PD-like neurochemical and behavioral outcomes. The optimized AAV2/rh10.TH.SNCA vector established a framework in which graded α-syn expression levels produced dissociable molecular, inflammatory, and behavioral phenotypes, ranging from early functional impairment to overt neurodegeneration. This now provides for a mouse model of PD where controlled nigral α-syn expression levels can capture distinct stages of disease progression. This is consistent with human data, where triplication of the wild-type α-synuclein gene alone can cause pathology, indicating that this model allows for use of a single reagent to determine the molecular, cellular and behavioral consequences of controlled increases in α-synuclein expression.

While AAV2/1 and AAV2/rh10 serotypes effectively transduced nigral neurons as seen previously ^33,36^, and the hSYN and TH promoters supported robust neuronal expression, our results demonstrate that efficient transduction alone is not sufficient to drive PD-relevant pathology or motor dysfunction. Among the promoters tested, hSYN achieved highest transduction efficiency in nigral neurons (AAV2/1.hSYN.SNCA), consistent with its known capacity for durable neuronal expression ^34,44,45^. Notably, AAV2/1.hSYN.SNCA failed to induce motor deficits and TH loss in both SN and striatum, whereas AAV2/rh10.TH.SNCA produced progressive motor impairment and nigral neurodegeneration despite more restricted transduction efficiency, especially when injected at low titer. These findings underscore the importance of cell-type specificity, local cellular context, and α-syn expression levels, rather than overall transduction rate alone, in determining pathological outcome.

Most of the AAV-based PD rodent studies report motor deficits when > 50% of SNc dopaminergic neurons are lost, which is consistent with clinical PD, but the severity and consistency of the resulting behavioral deficits have varied substantially across studies (**Table 1**). A few studies in rats suggest that a milder neuronal loss may be sufficient to induce molecular or motor changes. Notably, these studies were performed in females. We show that, high-titer AAV.TH.SNCA injections into young wild-type mice induced a significant ∼ 50% TH loss in both SN and striatum and motor deficits. Conversely, low-titer AAV.TH.SNCA(L) injections produced motor deficits and nigral neuropathological changes, without significant TH loss, consistent with early-stage PD pathology. Interestingly, lack of overt neurodegeneration was observed also in transgenic mouse lines overexpressing wild-type human α-syn under the rat TH promoter. A modest reduction in TH cell counts and motor deficits were observed only in aging conditions ^46–48^. These studies highlight the importance of characterizing other factors known to influence progression of human PD, including aging and sex, in future studies of variable expression levels with our AAV.TH.SNCA model.

The RNAscope *ISH* quantification of hSNCA transcripts at single-cell resolution demonstrated a roughly two-fold increase in hSNCA mRNA levels in nigral neurons transduced with AAV2/rh10.TH.SNCA vector injected at high-titer compared to low-titer. This supports a threshold model of α-syn toxicity in which sub-degenerative levels of α-syn induce pathological phosphorylation at Ser129, robust microglial activation, and sustained motor deficits in the absence of dopaminergic neuron loss. When this threshold is exceeded, achieved here through higher viral titers, significant TH+ neuronal degeneration emerges ^40,49,50^. This distinction is particularly important, as neuronal loss may obscure early molecular and inflammatory processes that are most relevant to disease initiation and progression. This also highlights how in some circumstances, relatively small changes in the number of AAV particles delivered to a target can lead to a corresponding linear change in transgene expression, with substantially different effects on neuronal responses to the transduced gene.

At sub-degenerative expression levels of α-syn, mediated by AAV2/rh10.TH.SNCA (L) injection, we observed a positive correlation between α-syn pSer129 abundance and Iba1+ microglial proliferation in the SN, which are both known pathological hallmarks of PD in clinical and animal studies ^13,18,40,51–55^. This supports the notion that microglial activation reflects a sensitive index of early neuropathological stress rather than a byproduct of overt neuronal death ^29,56,57^. Microglia are known to internalize and respond to extracellular α-syn, amplifying inflammatory cascades that may exacerbate synaptic and metabolic dysfunction^58,59^. These findings reinforce the role of neuroimmune interactions as early drivers of α-syn–mediated pathology.

Importantly, this threshold concept closely parallels human PD genetic cases. SNCA gene duplications and triplications in humans are associated with earlier disease onset, increased severity, and more rapid progression, directly implicating α-syn dosage as a determinant of pathology. Our AAV model recapitulates this dosage sensitivity experimentally, providing a translationally relevant system in which α-syn expression level - not merely its presence - dictates disease phenotype.

In conclusion, we generated a novel AAV vector with hSNCA transgene under the rat TH promoter, which achieves sustained transduction of nigral dopaminergic neurons and allows for distinct neurochemical and behavioral outcomes mimicking parkinsonian features. Our results demonstrate that precise modulation of α-syn expression is essential for revealing early pathogenic events that precede neuronal death. Lower expression levels enabled detection of early pathological markers, including α-syn hyperphosphorylation and microglial activation, while higher expression levels resulted in significant dopaminergic degeneration in young mice at eight months post-injection. This highlights how fine-tuning viral titer and promoter selection can model distinct stages of PD, from pre-degenerative dysfunction to overt neurodegeneration. Importantly, our findings show that neurochemical and behavioral alterations can occur independently of nigral cell loss, emphasizing that functional impairment in PD arises from molecular dysfunction prior to structural degeneration.

### Technical considerations and Limitations

Several limitations should be acknowledged. First, broad intra-group variability in transduction efficiency was observed for different AAVs, which is unavoidable with a direct injection model and is consistent with prior reports ^53,60^. Nevertheless, clear trends emerged that informed the selection of constructs for α-syn overexpression experiments. Second, ectopic mCherry expression from AAV.TH.mCherry in some TH negative cells in regions that do not contain DA neurons, may result from leaky TH promoter activity in precursor or non-dopaminergic populations ^61,62^. Mild motor changes observed over time in high-titer AAV.TH.mCherry controls highlight the importance of accounting for viral load effects when interpreting behavioral outcomes. While Ser129 phosphorylation is a widely accepted pathological marker, other post-translational modifications of α-syn (e.g., Ser87, Tyr39, Tyr125/133/136), which have also been reported to contribute to α-syn’s functional and pathological roles ^63,64^, were not assessed in this study and may further refine early pathological staging. Only two doses were tested here, so we do not know what the full linear range is with this vector in this brain region for viral particles injected to transgene expression, and additional titers within that range could lead to further differences in biological and behavioral effects. Finally, this study focused on male mice; future work should address potential sex-dependent differences in α-syn pathology and AAV transduction.

## CONCLUSIONS

Our data establish that wild-type human α-syn overexpression below a critical threshold induces neuroinflammation and α-syn hyperphosphorylation, resulting in motor deficits prior to dopaminergic neuron loss. When this threshold is exceeded, progressive neurodegeneration ensues, limiting the detection of early-stage molecular changes. The ability to induce graded α-syn pathology through controlled modulation and optimization of viral characteristics provides a unique opportunity to model both prodromal and degenerative stages of PD within the same framework. This novel, titrable AAV2/rh10.TH.SNCA mouse model provides a powerful and flexible platform to investigate the continuum of α-syn–driven pathology and neuroinflammation, and to test interventions aimed at halting PD progression at its early stages before neuronal loss occurs.

## METHODS

Additional details about Key Resources, such as reagents and antibodies, are included in the provided **Key Resource Table (Table 3)**.

**Table 3.**
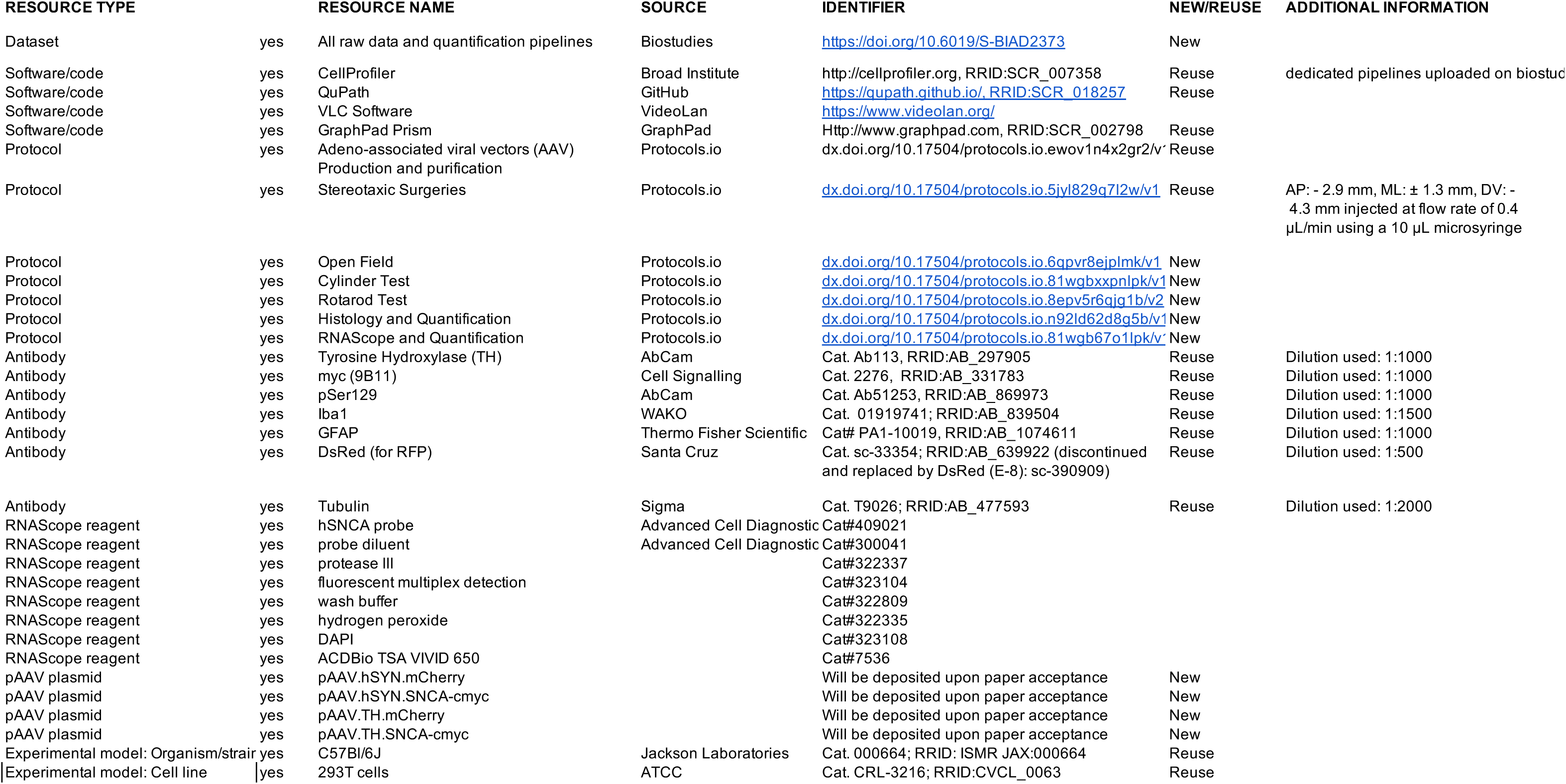
Key Resources.

### Generation of AAV constructs

We generated AAV vectors expressing mCherry encoding a red fluorescent protein (as a control) and the human wild-type α-syn transgene (SNCA) under the hybrid CMV enhancer/chicken β-actin (CBA) promoter ^33^, human Synapsin (hSYN) promoter ^34^ or a 2.5 kb rat Tyrosine Hydroxylase (TH) promoter ^35^ (**Figure 1a,b**). All the AAV plasmids used to generate AAV vectors were in a pAAV plasmid backbone including a Woodchuck Hepatitis Virus Posttranscriptional Regulatory Element (WPRE) sequence (**Figure 1a,b**).

The pAAV containing the mCherry transgene under CBA and hSYN promoters, and the pAAV.CBA.hSNCA with c-myc tag were previously cloned in the laboratory of Dr. Michael G. Kaplitt. The hSNCA-c-myc was subcloned into the pAAV.hSYN promoter plasmid within NcoI and HindIII restriction sites. The pAAV containing the rat TH promoter was purchased from Addgene (Plasmid #80336). The mCherry and hSNCA transgenes were excised from the pAAV.CBA.mCherry and pAAV.CBA.hSNCA-c-myc plasmids, respectively, and subcloned into the pAAV.TH promoter within XhoI and SalI restriction sites. Transgene expression from the plasmids was assessed via transient transduction (48 hours) in human embryonic kidney 293T (HEK-293T) cells (ATCC Cat. # CRL-3216) and western blotting (**Figure 1c**). Transgene expression from the packaged AAV vectors was validated *in vivo* via fluorescent immunolabeling (**Figure 1h**).

### Cell culture transfection and immunoblotting

HEK-293T cells were cultured in DMEM (Gibco), supplemented with 10% (vol/vol) FBS (Sigma-Aldrich) and 1% (vol/vol) penicillin-streptomycin (Gibco), at 37 °C in 95% humidified air and 5% CO2. Transient DNA transfection of our pAAV were performed using FuGENE6 transfection reagent (E2691) following manufacturer’s protocol (Promega Corp, Marison WI) in HEK-293T cells. Forty-eight hours following transfection, cells were lysed in lysis buffer (150mM NaCl, 50mM Tris pH 7.5, 2mM EDTA, 0.4%Triton-X, and protease and phosphatase inhibitor cocktail). Whole cell lysates were quantified by BCA assay and proteins were loaded on an SDS-PAGE and transferred to PVDF membrane. Because we were expecting a much lower expression of the transgenes from the plasmids containing the TH promoter compared to the plasmids with CBA promoter, to visualize protein expression levels in this cells system 50 μg of proteins were loaded on an SDS-PAGE for the pAAV.CBA plasmids and 100 μg for the pAAV.TH plasmids (**Figure 1c)**. Membranes were blocked with 5% milk in TBS-Tween (150 mM NaCl, 50 mM Tris at pH 7.5, and 0.1% Tween-20) and incubated with the primary antibodies overnight at 4°C. HRP-coupled secondary antibodies were incubated the next day at room temperature for one hour. Signal was detected with the Lumigen (Lumigen, Inc.) chemiluminescent reagents and images were taken with the VersaDoc MP 4000 using the Quantity One software (BioRad).

### Cell Culture and AAV Preparation

HEK-293T cells were cultured in DMEM (Gibco), supplemented with 10% (vol/vol) FBS (Sigma-Aldrich) and 1% (vol/vol) penicillin-streptomycin (Gibco), at 37 °C in 95% humidified air and 5% CO2.

The viral vectors used in this study expressed either human SNCA transgene or mCherry as a control. Two hybrid viral vector serotypes, AAV2/1 and AAV2/rh10, with AAV2 ITRs were used (**Figure 1a,b**).

Vector stocks were prepared by packaging the plasmids into mixed serotype AAV particles using a calcium phosphate transfection system as described previously ^33^. Cells were harvested and lysed at 72 h after transfection. The vectors were purified using iodixanol gradient and dialyzed against PBS with 2 mM MgCl2. AAV titers were determined by quantitative PCR (see also below).

### Quantitative Real Time PCR (qPCR)

To titer our purified viral vectors, AAV were processed as previously described ^33^ and viral genomes quantified via qPCR using FAST SYBR Green chemistry on the AB 7500 FAST Real Time PCR platform (Applied Biosystems) and primers to the WPRE element: WPRE-Fw: 5’-GGCTGTTGGGCACTGACAAT-3’; WPRE-Rev: 5’-CTTCTGCTACGTCCCTTCGG-3’. The relative number of full viral particles were calculated using the standard curve method by normalizing to known standard samples.

### Animals

Adult C5BL/6J male mice (4-months-old at time of AAV-injection, n=8-10 mice/group) (Jackson Laboratories; JAX stock # 000664) were kept on a reversed 12 h day-night cycle (11 am off/11 pm on) at a room temperature of 22 °C and 40–60% humidity. Animals received a standard chow diet and water ad libitum. All animal experiments were approved by the Weill Cornell Medicine Institutional Animal Care and Use Committees and performed according to the National Institutes of Health guidelines for the Care and Use of Laboratory Animals.

### Stereotactic injections

Mice were anesthetized by injecting a mixture of ketamine (110 mg/Kg) and xylazine (8 mg/Kg) intraperitoneally and were placed onto a stereotactic frame (Kopf Apparatus) with the skull flat between Lambda and Bregma. A small volume (0.1 ml) of local anesthetic agent bupivacaine (Marcaine 0.25%) was injected into the tissue adjacent to the intended skin incision line on the skull. The following coordinates were used for the injections into the SN (flat skull position): AP: - 2.9 mm, ML: ± 1.3 mm, DV: - 4.3 mm, below the dural surface as calculated relative to Bregma. Mice were injected with 1.5 μL of either AAV.mCherry (control vector) or AAV.SNCA into the right and left SN. AAVs were infused at a rate of 0.4 μL/min using a 10 μL micro syringe (World Precision Instruments) and an automated micro syringe pump (Kopf Apparatus). To allow for the diffusion of AAV into the tissue, the needle was left in place for 10 min after infusion and then slowly retracted at 0.2 mm/min. After the surgery, meloxicam (0.05 mg/Kg) was injected subcutaneously. Mice were kept warm in their cages until they fully recovered.

All AAV vectors were injected *in vivo* at a titer of 1×10^12^ vg/ml. AAV vectors containing the TH promoter were additionally injected at lower titer of 5×10^11^ vg/ml (designated TH(L)). AAV vectors with the CBA and hSYN promoters were injected at a titer of 1×10^12^ vg/ml. Vectors containing the TH promoter were injected either at lower titer of 5×10^11^ vg/ml (designated TH(L)).

### Behavioral tests

Mice were tested once per month over two consecutive weeks in the Open field arena, cylinder and rotarod (**Figure 2d**). The same investigator conducted behavioral tests. Testing occurred at the same time each day. As mice are nocturnal animals, behavioral testing took place during the dark phase of their daily cycle. Mice were habituated to the testing room for 2 hours before each session. Behavioral apparatuses were cleaned with 70% ethanol between trials.

#### Open Field test

The open field consisted of a squared plexiglass arena (40 cm x 40 cm) connected to an infrared beam system for locomotor tracking (Med Associates). Prior to recording of locomotor activity in the open field arena, mice were acclimated to the arena for 10 minutes over 2 consecutive days. On the third day, locomotion was recorded for 30 min and analyzed (Med. Associates). Spontaneous locomotor activity was measured as total traveled distance over 30 minutes (mean ± SEM).

#### Cylinder test

Mice were placed individually inside a clear glass cylinder (20 cm Ø, 30 cm height). The test started immediately and lasted 3 min. During the test session, mice were left undisturbed and were videotaped with a camera located at the front center of the cylinder. Mirrors were angled in the back and placed at the bottom to allow a 360 ° angle view. Wall contacts were analyzed by two independent operators blind to the experimental conditions, using a slow-motion video player (VLC software, VideoLan). The number of wall contacts with fully extended digits was counted and averaged (mean ± SEM).

#### Rotarod test

Mice were trained for two days to walk at an increasing speed (4.0 to 40 RPM) on a rotarod (Med Associates) for 5 consecutive trials per day, each lasting 5 min with an inter-trial time of 2-3 minutes. The latency to fall was automatically recorded by an infrared beam system. Coordination and balance on the rotarod were tested on the third day by averaging the latency to fall of the 5 trials (mean ± SEM).

### Brain fixation and histology

On completion of all behavioral assessments (∼8 months after AAV injections) mice were deeply anesthetized using sodium pentobarbital (50 mg/kg, i.p.) and intracardially perfused with cold heparinized PBS (30 mL) followed by 4% paraformaldehyde (PFA) (30 mL). Brains were rapidly removed and post-fixed in 4% PFA for 24 h and then transferred to a 30% sucrose solution for cryoprotection. Brains were sectioned into 40 μm thick coronal sections on a vibratome (VT100X Leica Biosystems, Buffalo Grove, IL) and stored in an anti-freeze solution (0.5 M phosphate buffer, 30% glycerol, 30% ethylene glycol) at – 20 °C. Sections were collected in 12 equally spaced series through the entire anterior-posterior extent of the SN and striatum and stored until further analysis.

### Immunofluorescent labeling of brain sections

Immunofluorescent labelling was performed on free-floating sections. For each histology experiment, 2 SN (-2.00 to -2.70 mm from Bregma) and 2 striatal (-2.90 to -3.50 mm from Bregma) sections per animal were used. Sections were washed three times with PBST buffer (0.2% Triton). To prevent bleed-through of the mCherry signal (AAV-mediated overexpression in control groups) into the wavelengths of the other fluorophores, sections were incubated for 6 hours at room temperature in a 3% H_2_O_2_ solution in PBST buffer (3 washes, 2 hours/each) prior to Immunofluorescent labelling. Afterwards sections were rinsed 3 times in PBST and blocking step was performed for an hour at room temperature. Primary antibody incubation was performed overnight at 4 °C. Blocking and primary antibody incubations were performed in 3% BSA in PBST (TH, Abcam, cat. Ab113, 1:1000; c-myc, Cell Signaling, cat. 2276 1:1000; Iba1, WAKO, cat. 01919741, 1:1000), or in 10% Donkey serum in PBST (pSer129, Abcam cat. ab51253, 1:1000). After the primary antibody overnight incubation, the tissue was rinsed in PBST and incubated with the respective AlexaFluor donkey secondary antibodies (all from ThermoFisher; anti-rabbit 488, cat. a21206, 1:1000; anti-sheep 647, cat. a21448, 1:1000: anti-rabbit 647, cat. 31573, 1:1000: anti-sheep 488, cat. A11015, 1:1000; anti-rat, 594 cat. a21209, 1:1000) for 60 min at room temperature in the specific blocking buffer. Sections were rinsed in PBST, mounted onto Superfrost Plus slides and coverslipped with ProLong diamond mounting medium (ThermoFisher).

### Immunofluorescent image acquisition and quantification

Histology images were acquired using a 10x objective mounted on an Olympus BX61TRF microscope and a Hamamatsu Digital Camera C13440 ORCA-Flash 4.0. To ensure unbiased data quantification, the analysis was performed by investigators blinded to experimental conditions. Estimates of total cell count (numbers of objects), and pixel density within regions of interest (ROI) in nigral and striatal sections were obtained using CellProfiler software ^65^. First, 10x images were acquired with an Olympus BX61TRF microscope, a Hamamatsu Digital Camera C13440 ORCA-Flash 4.0. The acquired 10x images were imported into the QuPath software ^66^. Then, regions of interest (ROIs) for the SN and striatum were drawn and exported in ImageJ. Afterward, quantification was performed with built-in-house pipelines in CellProfiler. Cells labeled for TH, mCherry, Iba1 and pSer129 were identified in a single-blinded setup based on cellular morphology and fluorescent intensity parameters. For each section the quantification was performed, and values from all the sections (counts/mm^2^) per each sample were averaged. Likewise, dedicated pipelines allowed the quantification of marker’s intensity per area (mm^2^) in the SN and striatum. Values from all the sections were averaged.

### Immunoperoxidase Histology and quantification

For each experiment, one SN and one striatal section per animal was selected and then punch coded in the cortex. Tissue sections from each experimental group were pooled into single containers to ensure identical reagent exposure ^67^.

SN and striatal sections from each condition were processed for TH. Sections were rinsed in 0.1M Tris-saline (TS; pH 7.6), blocked with 0.5% BSA in TS for 30 min, and incubated in sheep anti-TH (1:5000) diluted in 0.1% Triton-X and 0.1% BSA in TS for 24-hrs at room temperature followed by 24-hrs at 4°C. Next, sections were rinsed in TS and incubated in biotin-conjugated donkey anti-sheep IgG (Jackson ImmunoResearch Inc., West Grove, PA; RRID:AB_2340715) in 0.1% BSA and TS. Sections were washed in TS and incubated in Avidin Biotin Complex (ABC; Vectastain Elite kit, Vector Laboratories, Burlingame, CA) at half the manufacturer’s recommended dilution for 30-min. After rinsing in TS, the bound peroxidase was visualized by reaction in 3,3’-diaminobenzidine (Sigma-Aldrich, St. Louis, MO) and 0.003% hydrogen peroxide in TS for 10-minutes. All primary and secondary antibody incubations were carried out at 145 rpm, whereas rinses were at 90 rpm on a rotator shaker. Sections were mounted in 0.05 M PB onto gelatin-coated glass slides, dehydrated through an ascending series of alcohol to xylene, and coverslipped with DPX (Sigma-Aldrich).

To ensure unbiased data quantification, the analysis was performed by investigators blinded to experimental conditions. Quantification for TH labeling in the striatum was performed using previously established densitometric methods ^68–70^. Images were acquired using a Nikon Eclipse 80i microscope with a Micropublisher 5.0 digital camera (Q-imaging, BC, Canada) and IP Lab software (Scanalytics IPLab, RRID: SCR_002775). ImageJ64 software (Image J, RRID:SCR_003070) was used to measure the pixel density within regions of interest (ROI) in defined striatal regions. Background pixel density from non-labeled regions (e.g., anterior commissure) was subtracted to control for illumination variability and background labeling. Prior studies ^68^ demonstrated a strong correlation between pixel density and actual transmittance, confirming measurement accuracy.

### RNAscope *in-situ* hybridization (*ISH*)

The RNAscope *in situ* hybridization was used to quantify SNCA mRNA levels with fluorescent multiplex detection kit (#323104) following manufacturer instructions (Advanced Cell Diagnostics, Newark, CA, USA).

Nigral sections (30 μm) were washed twice in PBS 1x for 15 min and then mounted on Superfrost charged gold slides (Epredia) and let dry for 1 hour at RT. Afterwards, sections were rinsed in ddH_2_O, dried and then incubated for 1 hour at 60°C. Slides were then immersed in chilled 4% PFA diluted in PBS for 15 min. Next, were serially dipped in 50% and 70% EtOH diluted in H_2_O, and twice in 100% EtOH, and then incubated for 15 min at 60°C. Slides were incubated in hydrogen peroxide (#322335) for 10 min at RT, rinsed in ddH_2_O and then incubated for 15 min at 60°C. A hydrophobic barrier pen was then used to make a barrier around the perimeter of each section, and the barrier was completely dried at RT. The sections were then treated with Protease Ill (#322337) for 30 min at 40°C rinsed twice in PBS at RT, and then incubated with hSNCA probe-C1 (#409021) for 2 h at 40 °C. Next, the sections were rinsed twice in wash buffer (#322809) at RT and then successively treated with AMP 1 (30 min), AMP 2 (30 min), AMP 3 (15 min) reagents at 40°C. After each AMP treatment, the sections were rinsed twice in wash buffer at RT. After the last wash the sections were incubated with HRP-C1 (15 min), TSA Vivid™ Fluorophore 650 (Tocris Bioscience, part of Bio-Techne, # 7527) selected for C1 (1:2000) (30 min), and HRP-blocker (30 min) at 40°C. After each treatment the sections were rinsed twice in wash buffer at RT. After the last wash, sections were counterstained with DAPI (#323108) for 30 s at RT, completely dried at RT, and sealed with a coverslip using ProLong Gold Antifade reagent (Invitrogen) and then stored at 4 °C until imaged. Slides were imaged by within 48 h using an Olympus BX61TRF microscope, a Hamamatsu Digital Camera C13440 ORCA-Flash 4.0., and a 40x objective.

QuPath Software was used to quantify the number of transcripts for the hSNCA probe. Negative control images were generated by probing nigral sections with the RNAScope 4-plex negative control probe. A region of interest (ROI) was placed on the SN and the “Cell Detection” function was used to determine the number and position of cells in each ROI based on the DAPI nuclear stain (under the assumption that one nucleus represented one cell), and the ‘Subcellular Detection’ function was used to calculate the estimated number of transcripts for each target. Immunoreactive cells were identified as those with more than 3 estimated puncta.

An unspecific background signal was present in the AAV.mCherry samples due to the RNAscope ISH SNCA probe cross-reactivity with mouse SNCA mRNA. To correct for this, hSNCA transcript levels in AAV.SNCA-injected samples were normalized to the respective AAV.mCherry controls.

### Data analysis

Statistical analysis was performed with GraphPad Prism 9.0 (GraphPad Software) and significance was set at alpha < 0.05. Group comparisons were performed using ANOVA, with Tukey’s or Sidak’s correction for multiple comparison, or unpaired Student’s t-test, as stated in the Results section and figure legends. Graphs were generated in Prism 9 software. Data are presented as means ± SEM.

## ACKNOWLEDGMENTS

We thank the Neuroanatomy EM Core at Weill Cornell Medicine for assistance with immunocytochemistry experiments.

## AUTHOR CONTRIBUTIONS

Conceptualization: RM

Data curation: RM, SM, CRL, NDS, GS

Formal analysis: RM, SM

Funding acquisition: RM, MGK

Investigation: RM, SM, CRL, NDS, LGV, GS, WT, TAM

Methodology: RM, MGK, TAM

Project administration: RM

Resources: RM, MGK, TAM

Supervision: RM, TAM

Writing – original draft: SM, ERT, RM

Writing – review and editing: RM, SM, ERT, CRL, NDS, LGV, GS, WT, TAM, MGK

## DATA AVAILABILITY STATEMENT

All data are available in the main text or in the supplementary materials. For additional information and requests for resources and reagents corresponding author should be contacted, Roberta Marongiu.

