## Supplemental figures 1-4 for "A Versatile AAV-TH-SNCA Model to Study Early α-Synuclein Pathology and Intervention"

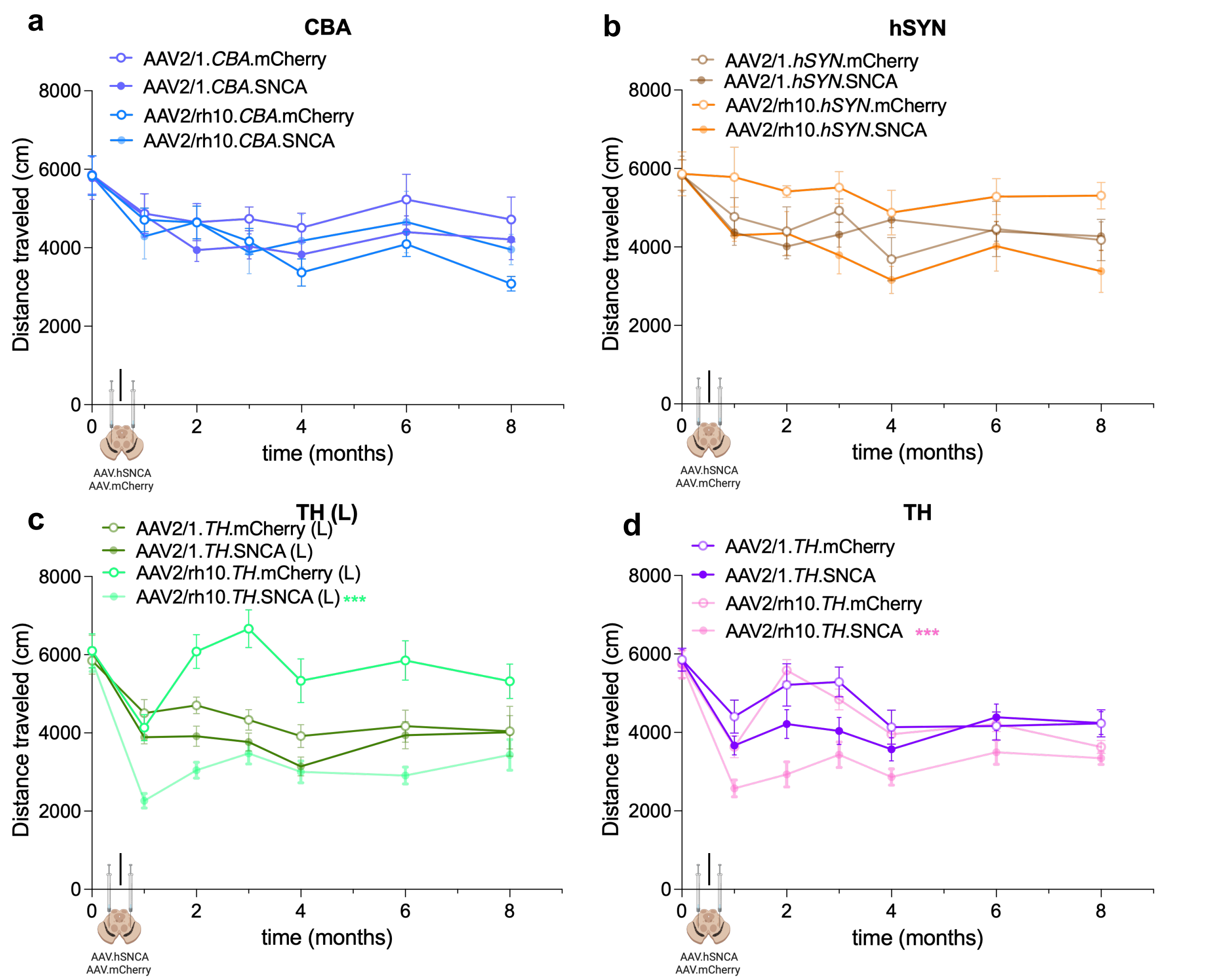


**Supplemental Figure 1. AAV2/rh10.TH.SNCA induce motor deficits in the open field test.** Results of the open field test are shown for mice injected with the AAV.mCherry and AAV.SNCA vectors carrying the CBA (a), hSYN (b) and TH promoter at the low (L) (c) or high (d) titer. Only mice injected with the AAV.TH.SNCA displayed deficits in the open field test. ***$p < 0.01$. AAV2/rh10.TH.SNCA (L): Two-way ANOVA, with Sidak's multiple comparison test, effect of time $F(3.47, 41.72)= 18.42$, ****$p < 0.0001$, effect of treatment $F(1, 13)= 34.96$, ****$p < 0.0001$, and effect of time*treatment $F(3.47, 41.72)= 6.694$, ***$p = 0.0005$. AAV2/rh10.TH.SNCA: Two-way ANOVA, with Sidak's multiple comparison test effect of time $F(3.89, 54.53)= 7.18$, ****$p = 0.0001$, effect of treatment $F(1, 14)= 18.89$, ***$p = 0.0007$, and effect of time*treatment $F(3.89, 54.53)= 7.18$, ***$p = 0.0001$). CBA, hybrid CMV enhancer/chicken β-actin promoter; hSYN, human Synapsin promoter; TH, Tyrosine Hydroxylase.

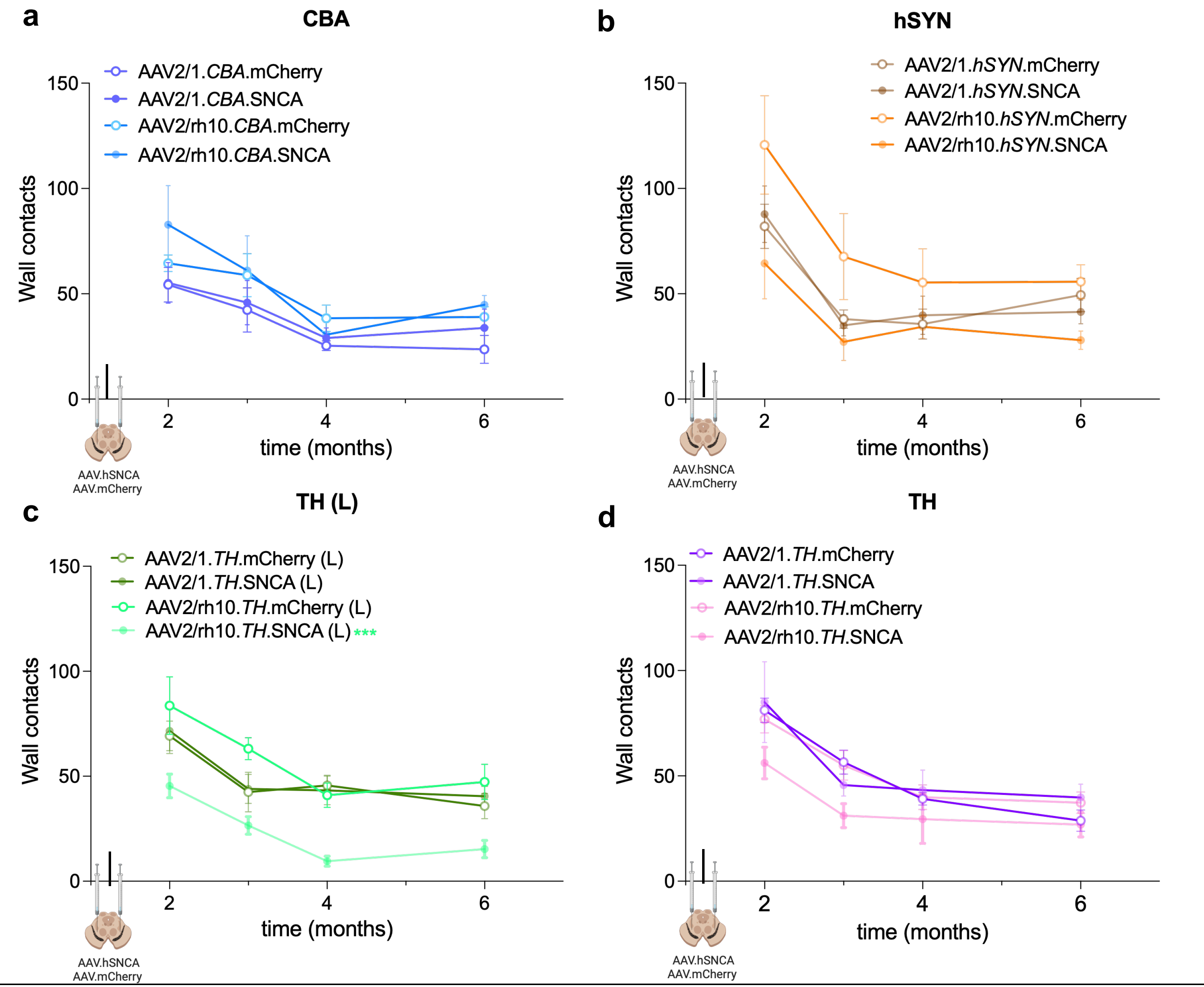


**Supplemental Figure 2. AAV2/rh10.TH.SNCA(L) induces motor deficits in the cylinder test.** Results of the cylinder test are shown for mice injected with the AAV.mCherry and AAV.SNCA with the CBA (a), hSYN (b), and TH promoter at the low (L) (c) and high (d) titer. Only mice injected with the AAV2/rh10.TH(L) showed deficits in the cylinder test. AAV2/rh10.TH.SNCA (L): Two-way ANOVA, with Sidak's multiple comparison test effect of time $F(1.58, 20.05) = 12.33$, ***$p = 0.0007$, effect of treatment $F(1, 13) = 37.38$, ****$p < 0.0001$, and effect of time*treatment $F(1.58, 20.05) = 0.104$, ns $p = 0.85$. CBA, hybrid CMV enhancer/chicken β-actin promoter; hSYN, human Synapsin promoter; TH, Tyrosine Hydroxylase.

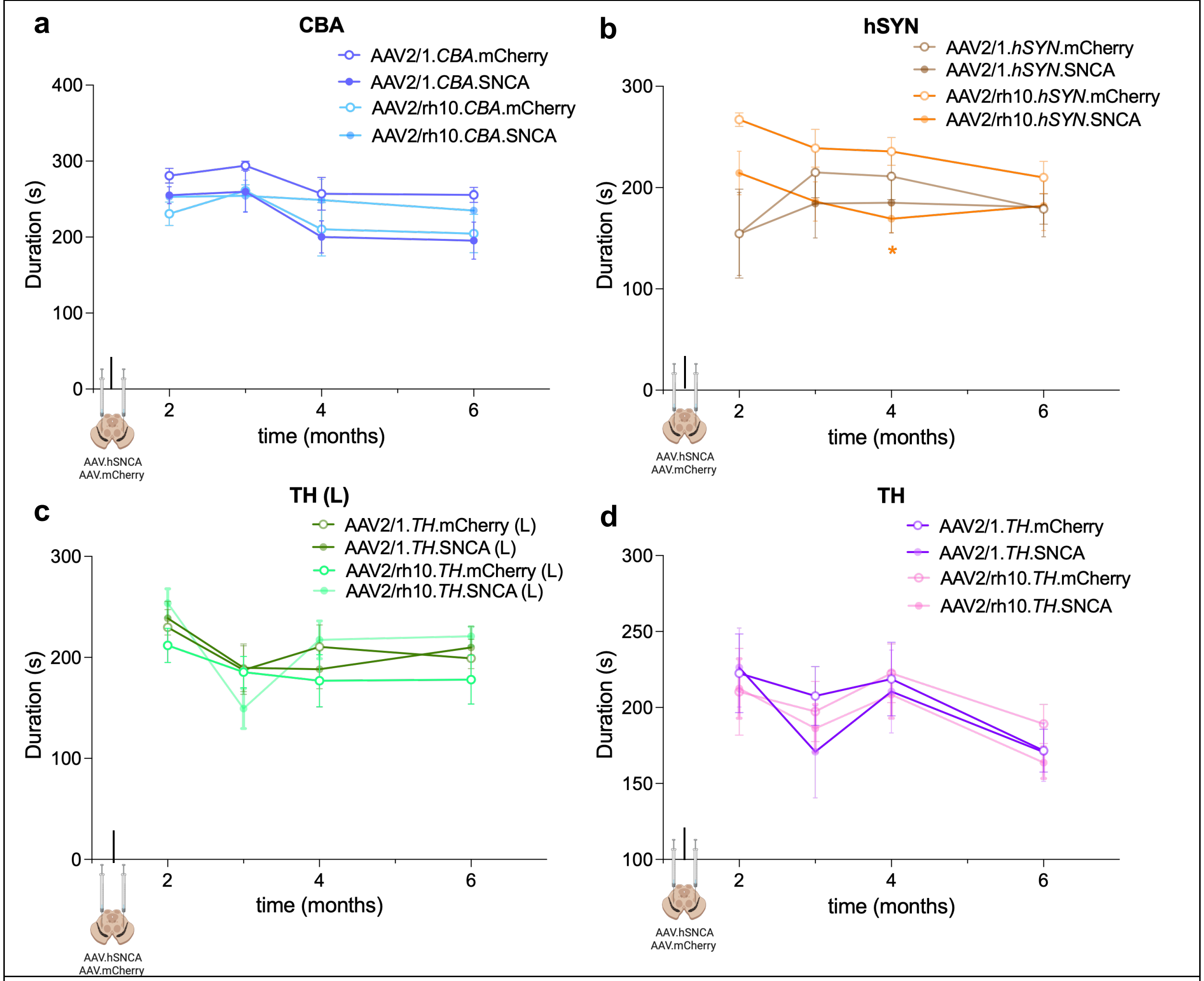


**Supplemental Figure 3. Motor deficits are not detected in the rotarod test.** Results of the rotarod test are shown for mice injected with the AAV.mCherry and AAV.SNCA with the CBA (a), hSYN (b), and TH promoter at the low (L) (c) and high (d) titer. No motor deficits were detected in the tested mice over time. Unpaired t-test *$p < 0.05$. Two-way ANOVA, Sidak's multiple comparison test effect of time $F(2.48, 29.84) = 5.46$, **$p = 0.006$, effect of treatment $F(1, 12) = 1.58$, ns $p = 0.23$, and effect of time*treatment $F(2.48, 29.84) = 2.89$, ns $p = 0.06$). CBA, hybrid CMV enhancer/chicken β-actin promoter; hSYN, human Synapsin promoter; TH, Tyrosine Hydroxylase.

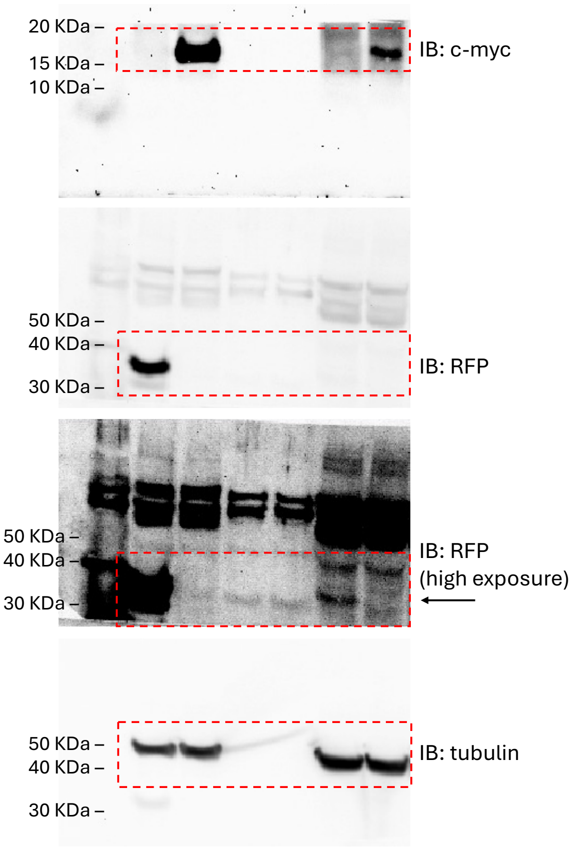


**Supplemental Figure 4.** Western blotting images used in Figure 1c.
